# Vaginal Community State Types (CSTs) Alter Environmental Cues and Production of the *Staphylococcus aureus* Toxic Shock Syndrome Toxin-1 (TSST-1)

**DOI:** 10.1101/2023.07.24.550353

**Authors:** Carla S. Maduta, John K. McCormick, Karine Dufresne

## Abstract

Menstrual toxic shock syndrome (mTSS) is a rare but life-threatening disease associated with use of high-absorbency tampons. The production of the *Staphylococcus aureus* toxic shock syndrome toxin-1 (TSST-1) is involved in nearly all cases of mTSS and is tightly controlled by regulators responding to the environment. In the prototypic mTSS strain *S. aureus* MN8, the major repressor of TSST-1 is the carbon catabolite protein A (CcpA), which responds to glucose concentrations in the vaginal tract. Healthy vaginal *Lactobacillus* species also depend on glucose for both growth and acidification of the vaginal environment through lactic acid production. We hypothesized that interactions between the vaginal microbiota (herein referred to as Community State Types, or CSTs) and MN8 depend on environmental cues, and that these interactions subsequently affect TSST-1 production. Using MN8 Δ1*ccpA* at various glucose levels, we demonstrate that the supernatants from different CSTs grown in vaginally defined media (VDM) significantly decrease *tst* expression. When co-culturing CST species with MN8 Δ*ccpA*, we show that *L. jensenii* completely inhibits TSST-1 production in conditions mimicking healthy menstruation or mTSS. Finally, we show that growing *S. aureus* in “unhealthy” or “transitional” CST supernatants results in higher IL-2 production from T cells. These findings suggest that dysbiotic CSTs may encourage TSST-1 production in the vaginal tract, and further indicates that the CSTs are likely important for the development of mTSS.

**IMPORTANCE:** In this study, we investigate the impact of the vaginal microbiota against *S. aureus* in conditions mimicking the vaginal environment at various stages of the menstrual cycle. We demonstrate that *L. jensenii* can inhibit TSST-1 production, suggesting the potential for probiotic activity in treating mTSS. On the other side of the spectrum, “unhealthy” or “transient” bacteria such as *G. vaginalis* and *L. iners* support more TSST-1 production by *S. aureus*, suggesting that CSTs are important in the development of mTSS. This study sets forward a model for examining contact-independent interactions between pathogenic bacteria and the vaginal microbiota. It also demonstrates the necessity of replicating the environment when studying one as dynamic as the vagina.

## INTRODUCTION

*Staphylococcus aureus* is a Gram-positive colonizer of human mucosal surfaces, including the vagina. This bacterium encodes a wide array of virulence factors, yet the superantigens are a unique subset of exotoxins which trigger an unconventional immune response. Superantigens force the activation of T cells through the cross-linking to MHC-II molecules, irrespective of peptide specificity of the T cell receptor (1). With up to 20% of the entire T cell repertoire potentially becoming activated (2), superantigens can initiate a highly pro-inflammatory state known as a cytokine storm which may lead to toxic shock syndrome (TSS). Staphylococcal TSS is a serious illness with symptoms including rash, high fever, hypotension, desquamation, and multiple organ dysfunction syndrome, which can become fatal if untreated (3).

The most concerning staphylococcal pathology in the vagina is the menstrual form of toxic shock syndrome (mTSS). The causative agent behind mTSS is the superantigen known as toxic shock syndrome 1 (TSST-1) (4). The production of TSST-1 by *S. aureus* is tightly controlled by a complex orchestration of regulators and environmental signals (5–7). Recently, it was discovered that the carbon catabolite protein A (CcpA) is the principle repressor of TSST-1 in the vaginal environment as a response to high glucose levels (8). Typically, the vaginal environment contains high glucose levels leading up to menstruation, at which point glucose concentration decreases significantly. The fluctuation in glucose level therefore likely contributes to the restriction of mTSS to the time-period of menstruation as a result of CcpA-mediated repression.

Other conditions which sustain the production of TSST-1 in this environment are high oxygen and carbon dioxide levels (9, 10), near neutral pH, and increased protein levels (11). Many of these conditions are met at menstruation, specifically with the use of high-absorbency tampons which can increase the oxygen levels in the typically anaerobic vagina (11, 12). As a result, the environmental cues present in the vaginal environment are critical for the restraint—or in the over production—of TSST-1.

The vaginal environment is strongly shaped by the microbiota. Classifications of the vaginal microbiota have been grouped into five Community State Types (CSTs), each fore fronted by a prominent bacterial species (13). *Lactobacillus crispatus*, *Lactobacillus gasseri*, *Lactobacillus iners*, and *Lactobacillus jensenii* dominate CST-I, CST-II, CST-III, and CST-V respectively, with CST-I and CST-III being the most common (13). Vaginal lactobacilli are often ascribed a protective function, with the exception of *L. iners* which is considered a transitional state (14). The final group, CST-IV, is a polymicrobial community composed mostly of strict anaerobes, as well as the prominent member *Gardnerella vaginalis* (15). CST-IV is strongly associated with dysbiosis and Bacterial Vaginosis (BV), the most common vaginal disorder among women of child-bearing age (16).

A protective role for vaginal lactobacilli is mainly attributed to their critical role in the acidification of the vaginal environment via the metabolism of glycogen into lactic acid. The appearance of lactobacilli in the vaginal microbiota at puberty, and their sustenance until menopause, is further postulated to be due to the role of estrogen in vaginal epithelial cell differentiation and release of glycogen (17, 18). For instance, the presence of *L. crispatus* has been directly correlated with estrogen levels (19) and use of hormonal therapy in menopausal women (20), and the use of estrogen rings among transgender men have restored *Lactobacillus* levels (21). However, the inability to explicitly quantify a relationship between estrogen and cell-free glycogen is hypothesized to be due to a lag time in the maturation of basal vaginal epithelial cells and the subsequent release of glycogen from superficial cells (22, 23). Nonetheless, free-glycogen is positively correlated with lactobacilli levels and low vaginal pH, with pre-menopausal women having higher levels of lactobacilli than post-menopausal women (24). The inverse relationship between glycogen and pH corresponds to the strong association between lactobacilli with both high levels of free glycogen and with low pH (22, 23). At menstruation, the decrease in glucose concentration results in a decrease in the abundance of lactobacilli, and an increase in microbial diversity, thereby increasing the vaginal pH (15).

Considering the production of TSST-1 is favoured in a low glucose environment with increased pH, the drop in estrogen and subsequent decrease in lactobacilli at menstruation creates conditions allowing for mTSS to occur. In this study, we aimed to investigate how CSTs I–V may inhibit or support *S. aureus* MN8 TSST-1 production in various *in vitro* conditions which mimic the vaginal environment. Using a *tst* transcriptional reporter and anti-TSST-1 immunoblot assays, we found that supernatants from representative CST spp. can significantly decrease TSST-1 expression in the absence of CcpA-mediated glucose repression. We further demonstrate that *L. jensenii* can overcome a variety of environmental stressors to inhibit the production of TSST-1 by *S. aureus* MN8 Δ*ccpA* and may be a rational probiotic candidate to pursue in order to prevent re-occurrence of mTSS.

## RESULTS

### Supernatants from CSTs I–V have repressive activity hidden behind glucose repression in VDM

Vaginally Defined Media (VDM) was developed and adapted to mimic the vaginal environment and its secretions, containing high glucose and glycogen concentrations (25, 26). Although VDM represents the conditions found in the vagina, one major environmental driver which is absent in the media is the signalling from the resident microbiota. To investigate the effects of CSTs I–V on the expression of *tst*, each CST representative was grown in VDM at various glucose concentrations and the supernatants were collected for use in a luciferase reporter assay. To account for the potential presence of potent inhibitory molecules in the supernatant and depletion of nutrients, the supernatants were diluted with their respective fresh media prior to the assays.

In supernatants containing 60mM of glucose, glucose-repression of *tst* was evident in the wild-type *S. aureus* MN8 reporter (Fig. 1A). Supernatant from *G. vaginalis* significantly decreased relative *tst* expression in wild-type MN8 with high glucose (Fig. 1A) with little influence on *S. aureus* growth (Fig. S1, Fig. S2). To relieve glucose repression, the assay was repeated with CST supernatants grown in VDM containing 0.7mM or 0mM glucose. Under these de-repressed conditions, most CST supernatants significantly decreased *tst* expression (Fig. 1B, Fig. 1C). As a second method of relieving glucose repression, the *ccpA* mutant reporter strain was used and similar trends compared with wild-type were observed. The CST supernatants all decreased *tst* transcription in high (60mM) and low (0.7mM) glucose (Fig. 1D, Fig. 1E); however, without glucose in the CST supernatants, there was no significant decrease except with the *L. iners* supernatant (Fig. 1F). Overall, these data indicate that different representative CST supernatants have repressive activity for TSST-1 transcription but that this is hidden behind CcpA-mediated glucose repression.

**Fig. 1.**
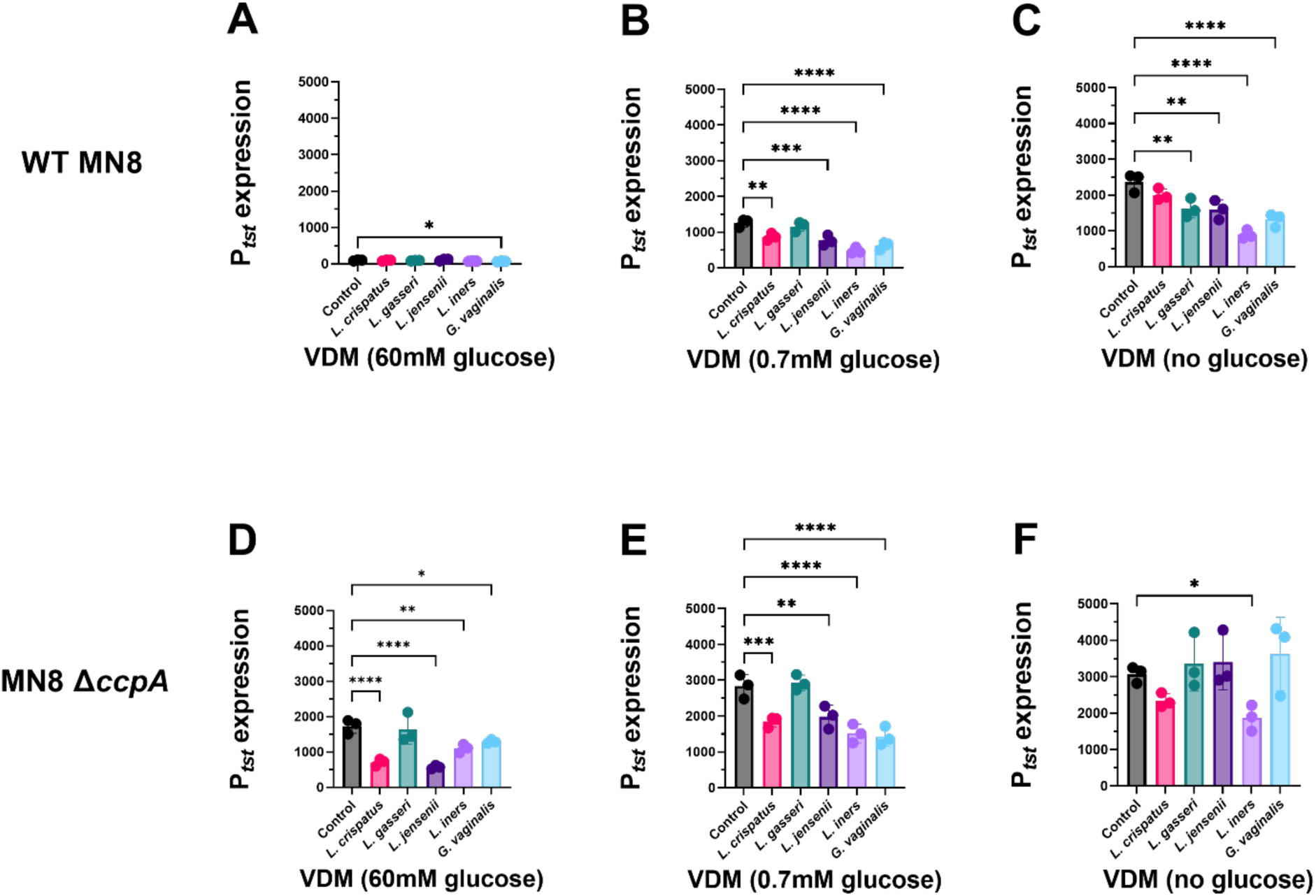
Supernatants from CSTs I–V have repressive activity hidden behind glucose repression in VDM. CSTs I-V were grown in standard VDM (60mM glucose), low glucose VDM (0.7mM glucose) or VDM lacking glucose (0mM glucose). CST supernatants were filtered (0.2 µm) and diluted with the respective fresh media at a 1/4 dilution. Wild-type MN8 (**A, B, C**) and MN8 Δ*ccpA* (**D, E, F**) containing the luminescence reporter plasmid pAmilux::P*_tst_* were then grown in the diluted supernatants. The assay was performed in the Synergy H4 reader at 37°C and continuous shaking on the medium setting. The relative luminescence units (RLUs) were standardized to the growth (OD_600_) (Figs. S1 and S2) to determine relative P*_tst_* expression. Error bars represent mean ± SD. Ordinary one-way Anova was performed with GraphPad Prism 9, no significance not shown (* P ≤ 0.05, ** P ≤ 0.01, *** P ≤ 0.001, **** P ≤ 0.0001).

### Supernatants from protective CSTs do not restrain production of the TSST-1 protein by MN8

While the luciferase reporter assay highlights repression of TSST-1 at the transcriptional level, the fully functional protein must be produced for mTSS to occur. To investigate whether “protective” CST supernatants (I, II, V) have similar activity at the transcriptional and translational levels, wild-type MN8 and MN8 Δ*ccpA* were grown in the diluted supernatants and TSST-1 production was assessed by Western blot. In control VDM or the associated CST supernatants, TSST-1 was not produced by wild-type MN8 (Fig. 2A). At 0.7mM and 0mM glucose levels, TSST-1 was produced across all conditions, highlighting the relief from glucose repression (Fig. 2B-C). Using the MN8 Δ*ccpA* strain, TSST-1 was detected in all supernatants regardless of glucose level, indicative again of the role of CcpA in glucose-mediated repression (Fig. 2D-F). Altogether, these results indicate that the restriction of *tst* by the CST supernatants does not abrogate production of the protein.

**Fig. 2.**
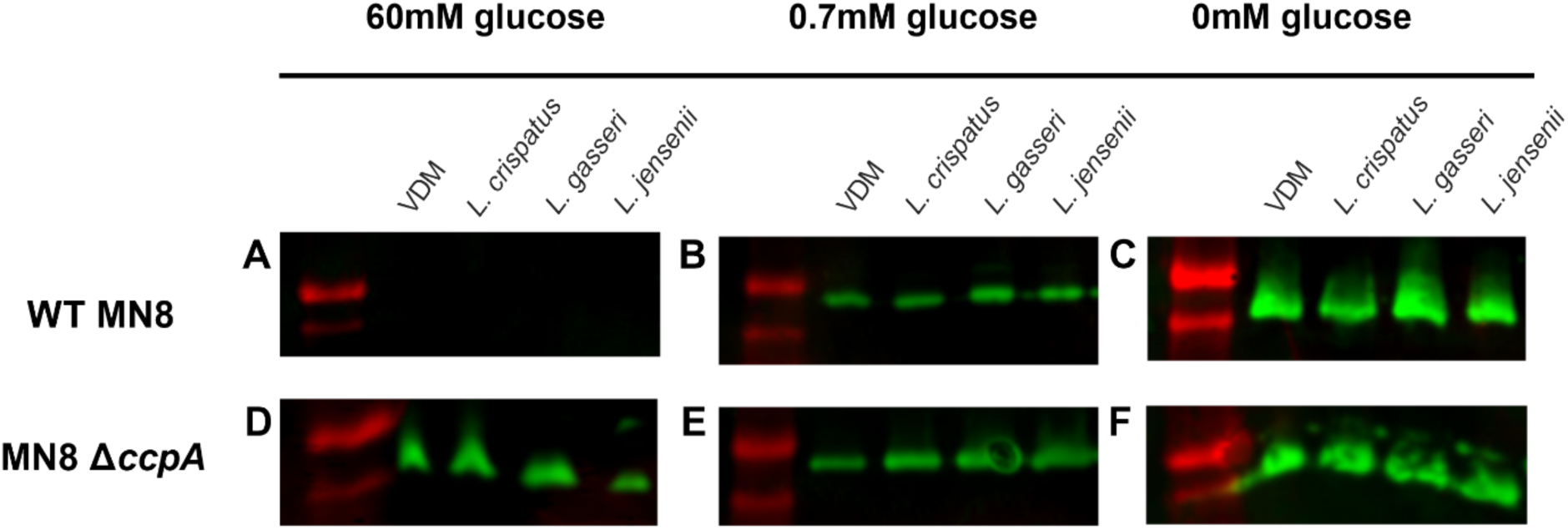
Supernatants from CSTs I, II and V do not eliminate production of TSST-1. CSTs I, II and V were grown in standard VDM (60mM glucose), low glucose VDM (0.7mM glucose) or VDM lacking glucose (0mM glucose). CST supernatants were filtered (0.2 µm) and diluted with the respective fresh media at a 1/4 dilution. Wild-type MN8 (**A, B, C**) and MN8 Δ*ccpA* (**D, E, F**) were then grown in the diluted supernatants. Supernatants were TCA precipitated (standardized to 12 OD_600_ units), followed by SDS-PAGE and anti-TSST-1 Western blotting.

### Supernatants from transitional and dysbiotic CSTs may increase production of TSST-1

To examine how transitional or dysbiotic CST supernatants (III, IV) affect TSST-1 production, supernatant experiments were performed using *L. iners* and *G. vaginalis*, respectively. In standard VDM (60mM glucose), TSST-1 was not produced by wild-type MN8, although production was recovered as glucose levels decreased (Figs. 3A-C). Using the MN8 Δ*ccpA* mutant, TSST-1 was produced across all glucose conditions in both fresh VDM and CST supernatants (Figs. 3D-F). In 0.7mM glucose VDM conditions, the TSST-1 bands from MN8 and MN8 Δ*ccpA* grown in *L. iners* and *G. vaginalis* supernatants appeared more intense relative to the VDM control (Fig. 3B, 3E), overall suggesting that TSST-1 production may be augmented in the presence of *L. iners* and *G. vaginalis* supernatants.

**Fig. 3.**
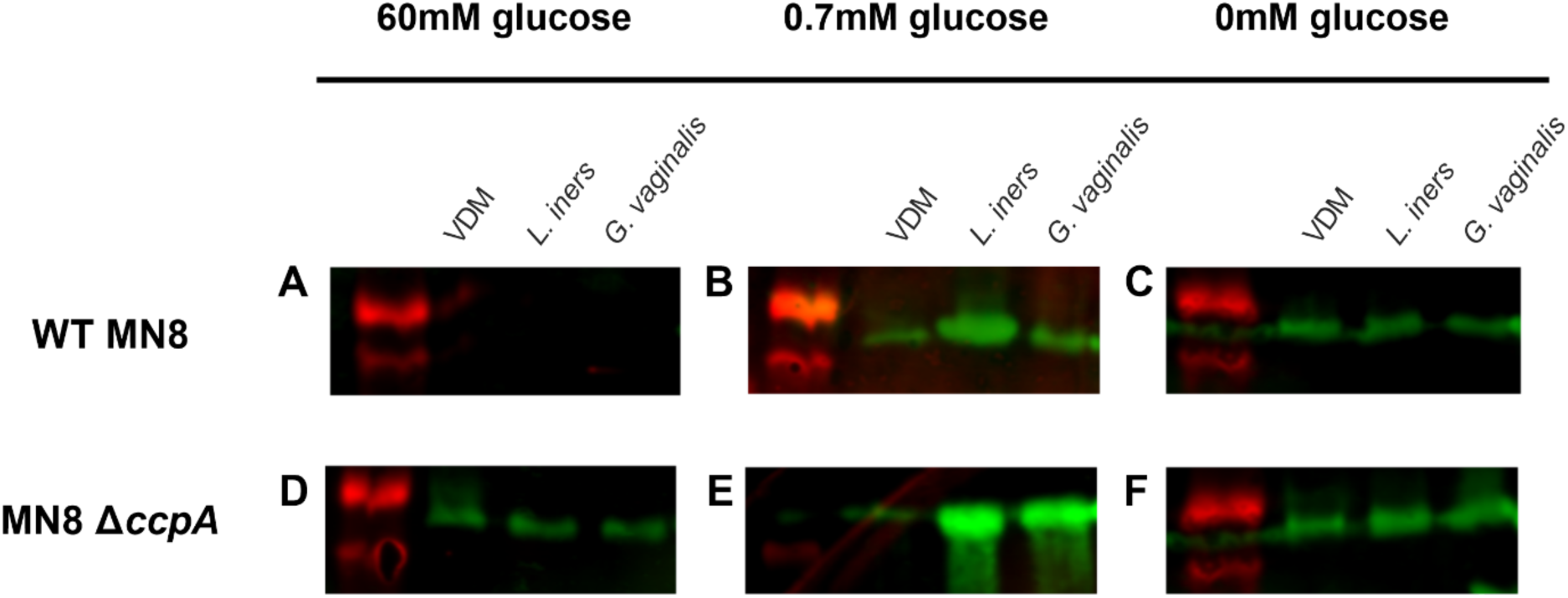
Supernatants from CSTs III and IV may augment production of the TSST-1 protein. CSTs III and IV were grown in standard VDM (60mM glucose), low glucose VDM (0.7mM glucose) or VDM lacking glucose (0mM glucose). CST supernatants were filtered (0.2 µm) and diluted with the respective fresh media at a 1/4 dilution. Wild-type MN8 (**A, B, C**) and MN8 Δ*ccpA* (**D, E, F**) were then grown in the diluted supernatants. Supernatants were TCA precipitated (standardized to 12 OD_600_ units), followed by SDS-PAGE and anti-TSST-1 Western blotting.

### Transitional and dysbiotic CST supernatants are capable of increasing T cell activation by S. aureus

Although CST supernatants downregulated *tst* at the transcriptional level during the exponential growth phase (Fig. 1), evaluating TSST-1 protein at stationary phase demonstrated that this exotoxin continued to be produced (Fig. 2, Fig. 3). This suggests that although the microbiota can have restricting effects on TSST-1 production, once the environmental conditions for TSST-1 production are met, the toxin will be produced. This led us to inquire whether the CST supernatants could alter the T-cell activation capacity of *S. aureus*. We exposed peripheral blood mononuclear cells (PBMCs) to the various titrated *S. aureus* supernatants to evaluate T cell responses. To ensure toxin production, the assay was performed using samples from experiments at 0.7mM glucose VDM with aeration to mimic environmental conditions typical of mTSS. With wild-type MN8 and the *ccpA* mutant, all supernatants from *S. aureus* grown in “healthy” CST supernatants were capable of inducing dose-dependent IL-2 production similar to *S. aureus* grown in the VDM control media (Fig. 4). However, *S. aureus* grown in *L. iners* and *G. vaginalis* supernatants were able to induce peak IL-2 concentrations at ~10-fold higher dilution, indicating stronger potency (Fig. 4). Among the blood donors, all displayed the same trends in T cell activation, however one donor was more sensitive to *S. aureus* exposed to *L. iners* and *G. vaginalis* supernatants. These results suggest an interesting trend that supernatants from transitional or dysbiotic CSTs may promote stimulation of T cells by *S. aureus*.

**Fig. 4.**
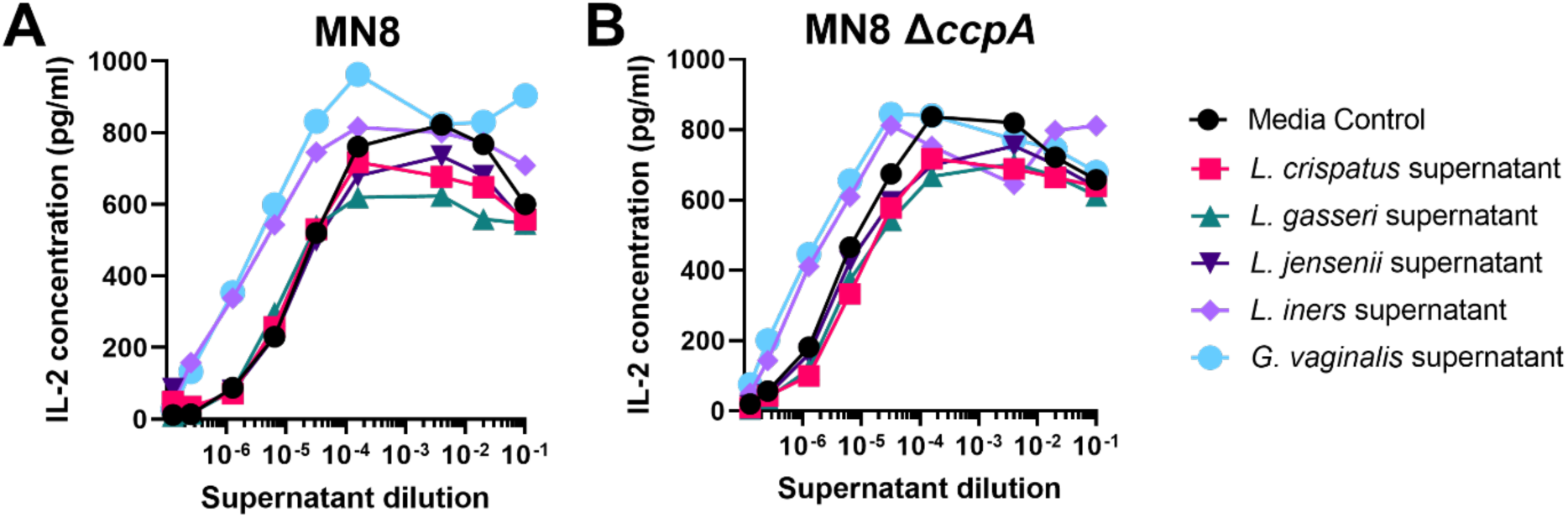
Transitional and dysbiotic CST supernatants enhance activation of T cells by *S. aureus* MN8. Representative CST species were grown in VDM 0.7mM glucose. CST supernatants were filtered (0.2 µm) and diluted with the respective fresh media at a 1/4 dilution. Wild-type MN8 (**A**) and MN8 Δ*ccpA* (**B**) were grown in the diluted supernatants with aeration, to mimic mTSS. The *S. aureus* supernatant was filtered and added to human PBMCs for 18h. Cell supernatants from PBMCs were assayed for IL-2 by ELISA as a measure of T cell activation. Data represent the mean from three healthy donors.

### Lactobacilli in co-culture with S. aureus MN8 lacking CcpA repression can inhibit production of TSST-1 in conditions mimicking healthy menstruation and menstrual toxic shock syndrome

The luciferase transcriptional reporter assays and supernatant experiments examined regulation of TSST-1 by growing both actors independently of each other; however, the vagina harbours a multitude of bacterial species simultaneously. These bacterial species may have interactions dependent on signalling that only occurs when both species are present, as opposed to solely the bacterial products found in spent media. In order to investigate the production of TSST-1 in a context to better resemble interactions that take place within the vagina, *S. aureus* MN8 was co-cultured with each of the protective CST representatives. Co-cultures were performed with *S. aureus* on one side of a 0.65 µm membrane, and one of the lactobacilli on the other, to allow for contact-independent signaling (Fig. S3). Each side of the co-culture contained VDM with various glucose concentrations, and experiments were performed aerobically to mimic mTSS and microaerophilically to mimic the otherwise low oxygen state of the vagina. Transitional and dysbiotic CSTs were excluded from co-culture experiments given the evidence suggesting they may promote TSST-1 production (Fig. 4).

In co-cultures performed with aeration, wild-type MN8 did not produce TSST-1 at 60mM glucose (Fig. 5A), yet this repression was relieved at 0.7mM glucose (Fig. 5E) and 0mM glucose (Fig. 5I). Under microaerophilic conditions, TSST-1 was not produced at 60mM (Fig. 5B) or 0.7 mM glucose (Fig. 5F), but without glucose TSST-1 was faintly detectable in co-cultures with *L. crispatus* and *L. gasseri* (Fig. 5J).

**Fig. 5.**
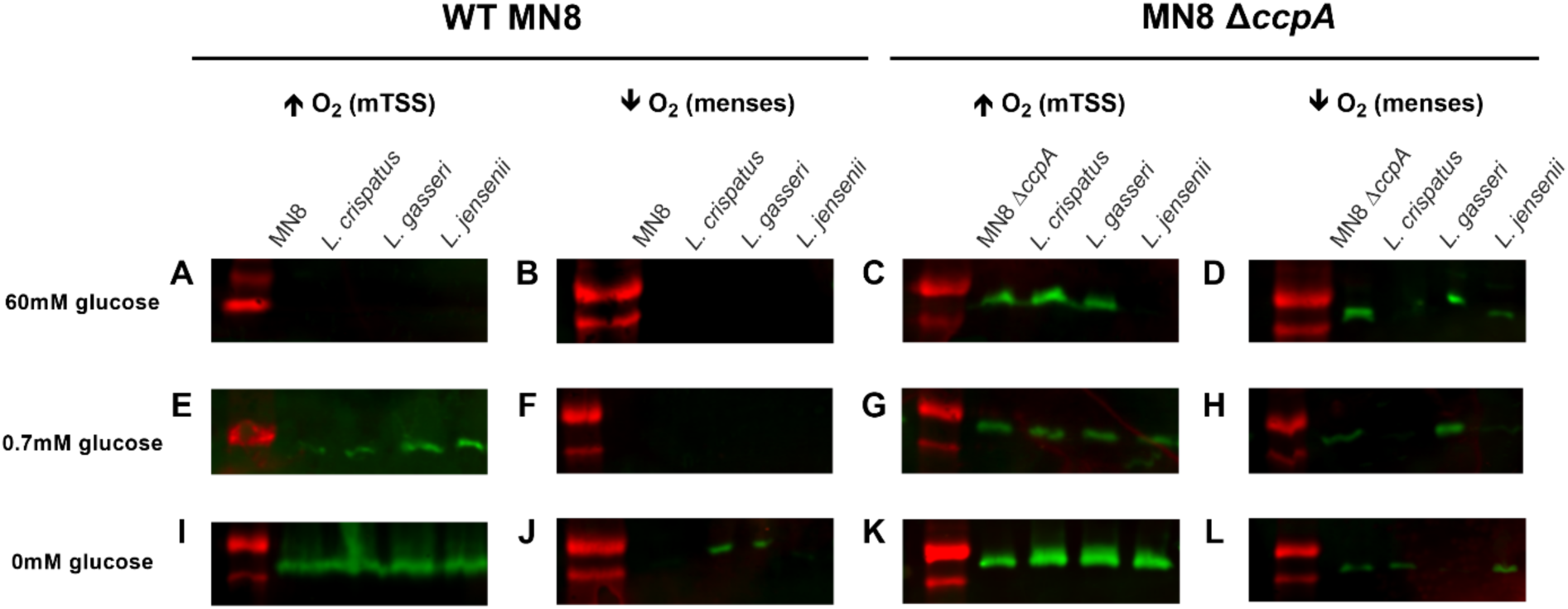
Environmental conditions affect regulation of TSST-1 by healthy lactobacilli co-cultured with *S. aureus* MN8. A co-culture apparatus separated two bacterial species with a 0.65 µm filter, with each side containing 20 mL of media, was used. Lactobacilli representing CSTs I, II and V were co-cultured with either wild-type MN8 or MN8 Δ*ccpA* in VDM at the indicated glucose concentrations. Control co-cultures were performed with the same *S. aureus* strain on either side of the membrane. Aerobic co-cultures to mimic mTSS environmental conditions were performed at 250 rpm and 37°C (**A, E, I, C, G, K**), while microaerophilic co-cultures to mimic healthy menstruation environmental conditions were performed without shaking at 37°C (**B, F, J, D, H, L**). TCA precipitations were performed the following day, standardized to 12 OD_600_ units, followed by SDS-PAGE and anti-TSST-1 Western blotting.

Using the MN8 Δ*ccpA* mutant in aerobic conditions, TSST-1 was produced in all co-cultures, except with *L. jensenii* at 60mM glucose (Fig. 5C). When glucose was reduced, this repressive activity by *L. jensenii* was lost (Fig. 5G, Fig. 5K). In microaerophilic co-cultures, *L. jensenii* maintained its repressive activity at both 60mM (Fig. 5D) and 0.7mM glucose (Fig. 5H), while *L. crispatus* gained this activity relative to the aerobic co-cultures (Fig. 5C, Fig. 5G). At 0 mM glucose (Fig. 5L), a switch between repressive activity was seen among the lactobacilli, as *L. crispatus* and *L. jensenii* co-cultures supported toxin production, while the *L. gasseri* co-culture did not.

Given that *L. jensenii* strongly restricted the production of TSST-1 in co-culture conditions that would mimic mTSS, we performed quantitative anti-TSST-1 ELISAs to confirm the qualitative Western blot analyses. In conditions with high aeration, co-cultures containing MN8 *ΔccpA* on either side of the membrane produced relatively high amounts of TSST-1 as average TSST-1 production at both 0.7mM and 60mM glucose was ~1500 ng/mL (Fig 6A, 6C). However, TSST-1 was essentially undetectable using this assay from aerobic co-cultures with MN8 *ΔccpA* and *L. jensenii* (Fig 6A, 6C). In contrast, microaerophilic co-cultures had limited TSST-1 production in all conditions, indicative of the strong repression that occurs when oxygen is limited, likely due to the regulator SrrAB (Fig 6B, 6D) (9). Overall, these results reveal that *L. jensenii* is able to significantly inhibit TSST-1 production in conditions mimicking mTSS when glucose-repression is relieved.

**Fig. 6.**
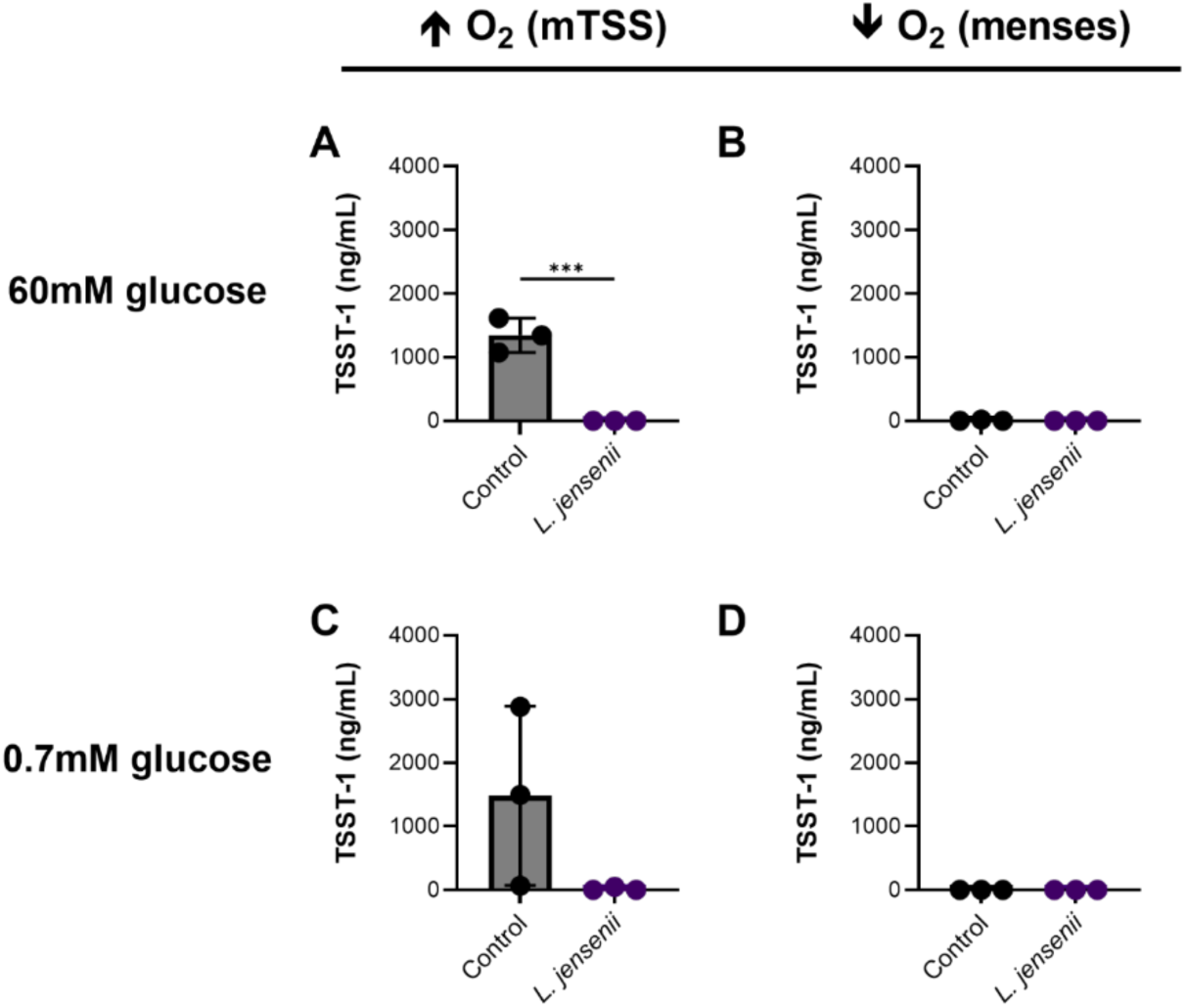
*L. jensenii* in co-culture with *S. aureus* MN8 Δ*ccpA* inhibits production of TSST-1. A co-culture apparatus separated MN8Δ*ccpA* from *L. jensenii* with a 0.65 µm filter, with each side containing 20 mL of VDM. Control co-cultures were also performed with MN8 Δ*ccpA* on either side of the membrane. Aerobic co-cultures which mimic mTSS were performed at 250 rpm and 37°C (**A, C**), while microaerophilic co-cultures which mimic healthy menstruation were performed without shaking at 37°C (**B, D**). *S. aureus* supernatants were collected the following day, filter-sterilized, and stored at −20°C until used in a quantitative TSST-1 ELISA. Unpaired t-tests were performed using GraphPad Prism 9, no significance not shown (*** P ≤ 0.001).

### Aeration is a determining factor in *S. aureus* virulence factor modulation by *L. jensenii*

The finding that *L. jensenii* could dramatically decrease production of TSST-1 in the absence of glucose repression led us to perform transcriptomic analysis of these notable co-cultures. For these experiments, we compared transcriptional profiles of *S. aureus* Δ*ccpA* grown on both sides of the co-culture apparatus with *S. aureus* Δ*ccpA* grown with *L. jensenii* for 4 hours post-inoculation. We found that co-cultures mimicking mTSS—containing 0.7mM glucose and high aeration—had 404 downregulated genes and 429 upregulated genes for a total of 833 genes differentially expressed. Of these genes, not only was there a downregulation of *tst* expression, there was an overall downregulation of multiple secreted virulence factors including PSMα genes, γ-hemolysin and leukocidins (Fig. 7). Notable genes which were upregulated were a variety of nitrate metabolism genes, sugar phosphotransferase systems, as well as the superantigen-like protein 1 (SSL1). In contrast, co-cultures with no aeration, which mimic healthy menstruation, had only 6 genes differentially expressed, and did not include *tst* (Fig. S4). Overall, this analysis suggests that the transcriptional profile of *S. aureus* MN8 Δ*ccpA* can be dramatically altered to a ‘less virulent’ state within as little as 4 hours of exposure to *L. jensenii* in an environment that would otherwise support mTSS.

**Fig. 7.**
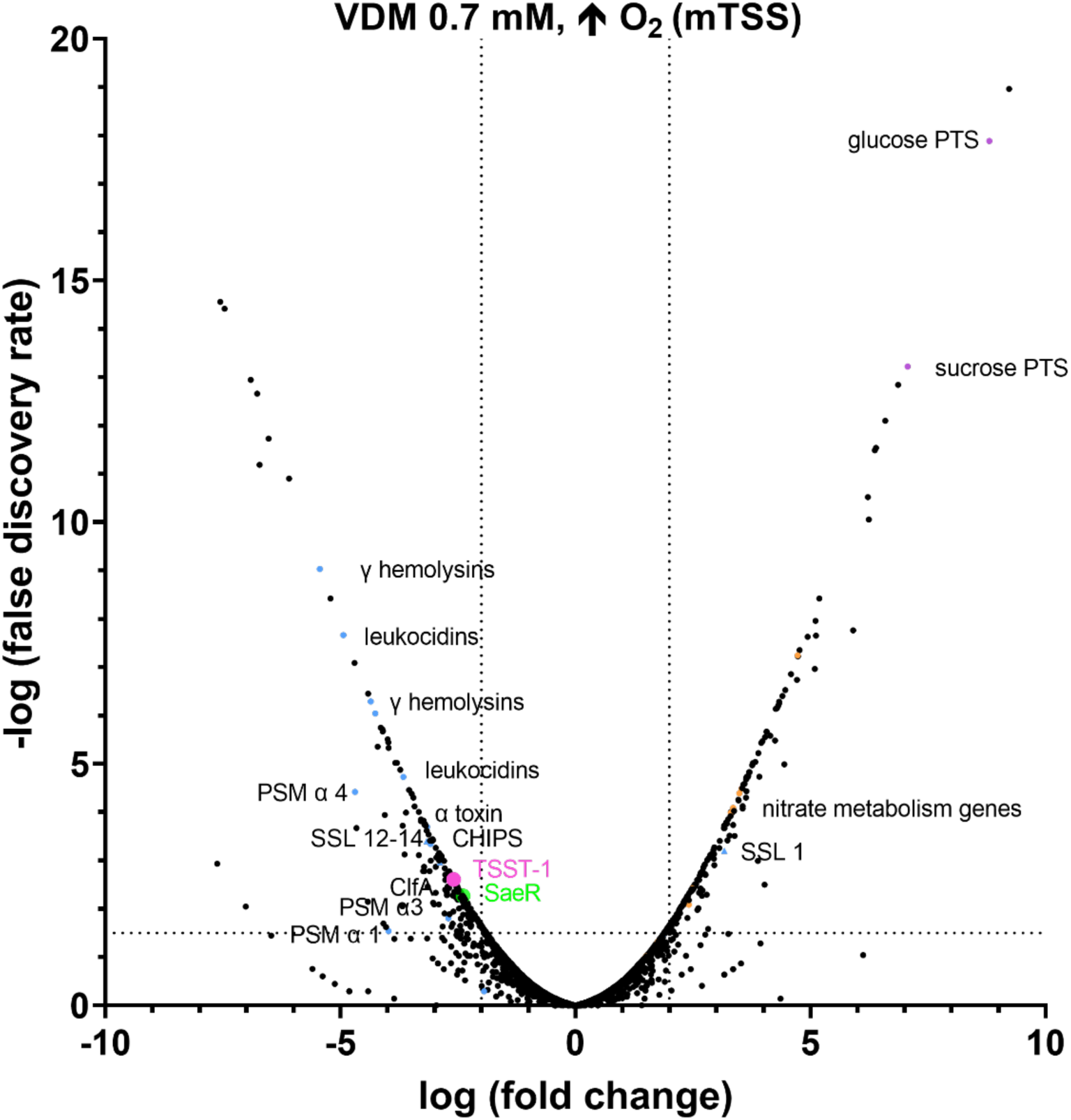
*L. jensenii* in co-culture with *S. aureus* MN8 Δ*ccpA* downregulates virulence factor expression in conditions mimicking mTSS. A co-culture apparatus separated MN8 Δ*ccpA* from *L. jensenii* with a 0.65 µm filter, with each side containing 20 mL of VDM 0.7mM glucose. Control co-cultures were also performed with MN8 Δ*ccpA* on either side of the membrane. Co-cultures were performed aerobically at 250 rpm and 37°C to mimic mTSS. Each dot in the volcano plot represents a single transcript, and those above the horizontal threshold indicate P < 0.001, while vertical thresholds indicate log fold changes greater than 2 in the presence of *L. jensenii*.

## DISCUSSION

The role of environmental cues in the regulation of the superantigen TSST-1 has been well established *in vitro* (8, 9, 27, 28). Through a wide range of repressive signals, expression of TSST-1 appears to be highly controlled by the causative bacterium *S. aureus* such that mTSS can occur only in very specific conditions. In this study, we have further established that the contributions of CSTs I–V are likely to also play a role in the prevention, or exacerbation, of mTSS.

While we previously established the importance of glucose concentration as a key environmental determinant in TSST-1 expression (8), we aimed to revisit the role of glucose in the context of the vaginal microbiota. At puberty, the increase in estrogen is believed to coincide with the accumulation of glycogen in vaginal epithelial cells, and the subsequent appearance of lactobacilli in the vaginal environment (17). Through the use of VDM at various glucose concentrations, we confirmed the importance of CcpA in repressing *tst* transcription (7, 8), as expression from the promoter and production of the protein increased in the absence of glucose (Fig. 1C, Fig. 5I) or in the MN8 Δ*ccpA* mutant (Fig. 1D, Fig. 5C). Given that *L. crispatus*, *L. gasseri*, and *L. jensenii* restrict the production of TSST-1 both in the presence and absence of glucose in media (Fig. 5C, 5D, 5H, 5L), this would strongly suggest that glycogen in the media is being catabolised in sufficient quantities to support lactobacilli growth. Although glycogen is critical for growth of lactobacilli and acidification of the vaginal environment, most lactobacilli species do not encode a known enzyme capable of degrading glycogen (29–32). Instead, these lactobacilli must rely on amylases of human and other microbial origins to catabolize the polymer into smaller sugars such as glucose and maltose, which can then be used to produce lactic acid (33, 34). While *L. crispatus* isolates have been identified to encode a pullulanase which sustains the growth of the bacteria on glycogen (31, 32), many strains contain non-functional copies. Although the strain used in this study does not contain these deletions, it demonstrated reduced growth in the absence of glucose (Fig. S5A) that was nearly identical to *L. jensenii* which does not encode a pullulanase (Fig. S5C). This suggests the *L. crispatus* pullulanase is non-functional in the conditions tested. However, an α-amylase has been previously characterized and expressed from *S. aureus* ATCC12600 (35), and we believe this gene is also expressed in *S. aureus* MN8. This would indicate that the small sugars released would be able to cross the co-culture membrane and support the growth of the lactobacilli (Fig. 5).

Likewise, a protective role of lactobacilli in the vaginal environment has been attributed to the decrease in vaginal pH through the production of L- and D-lactic acid via glycogen and glucose metabolism (22, 36–38). Given the significant reductions in *tst* expression demonstrated in the luciferase assays (Fig. 1), on the surface this would suggest that a decrease in pH may be the mechanism of action through which lactobacilli both restrict *S. aureus* growth, and also limit *tst* expression in the vaginal niche. However, these protective lactobacilli are all capable of decreasing the pH of VDM to similar values, yet throughout the study we show inter-species differences in the restriction of TSST-1 production. Recently, Schlievert et al. (2023) found that the probiotics *Lactobacillus acidophilus* LA-14 and *Lacticaseibacillus rhamnosus* HN001 can inhibit TSST-1 production, with the latter doing so in a pH dependent manner via the SrrA/B two-component system (39). This activity appears to be distinct from *L. jensenii* as *srrAB* expression was not affected in our transcriptome analysis. It may be possible that different D-/L-lactic acid ratios play a role in the interspecies differential regulation of *tst*, as *L. jensenii* is the sole lactobacilli in this study that only produces the D isomer, which has been implicated in protective effects relative to the L isomer (40, 41). Nonetheless, luciferase assays demonstrate decreased *tst* expression even in conditions of low or no glucose (Fig. 1B, 1C, 1E, 1F), which are environments that do not support sufficient acidification via lactic acid production. Ultimately this indicates that pH is not likely the principal method of repression and some, if not all, lactobacilli used in our study have additional mechanisms through which TSST-1 production is restrained. For example, it was found that the vaginal isolate *L. reuteri* RC-14 produces cyclic dipeptides which act in Agr dependent and independent manners to repress *tst* (42); however, it is unclear whether the lactobacilli in this study—or any other species and strains—produce these dipeptides.

One of the most striking results from this study is the ability of *L. jensenii*, as well as *L. crispatus*, to nearly entirely eliminate the production of the toxin in co-cultures with MN8 already lacking repression from CcpA and glucose (Fig. 5C, Fig. 5D, Fig. 5H, Fig. 6). This indicates that the TSST-1 repressive mechanisms employed by lactobacilli can be ‘hidden’ behind glucose repression and may function as a ‘backup’ defense mechanism. Conditions where this repression by lactobacilli occurred were mostly microaerophilic, which is consistent with knowledge that lactobacilli strive in low oxygen environments. However, given that *L. jensenii* completely inhibited TSST-1 production in a highly aerobic environment, this may indicate it is the most protective lactobacilli as it can overcome environmental stressors (Fig. 5C, Fig. 6A, 6C). Of note is the ability of *L. jensenii* to drastically reduce the production of TSST-1 in a contact-independent manner, as opposed to *L. jensenii* RC-28 which required physical contact with *S. aureus* MN8 (43). This indicates there must be key genomic differences between *L. jensenii* RC-28 and ATCC25258, which confer different protective functions in the context of mTSS. Overall, the co-cultures demonstrate that *L. jensenii* offers protection in the most environmental conditions compared to the other CST lactobacilli and merits further investigation as a potential probiotic candidate, specifically for those who have experienced mTSS in order to prevent re-occurrence.

The transcriptomic analysis revealed that most known *tst* regulators were not differently expressed in the presence of *L. jensenii* (Fig. 7A), with the exception of the major activator SaeR. Given that activation from SaeRS is required for TSST-1 expression (5), this provides us with an indication of the pathway that is targeted by *L. jensenii* in order to limit TSST-1 production. The decrease in *saeR* expression is also consistent with a downregulation of numerous virulence factors such as α-hemolysin, CHIPS, γ-hemolysin, and leukocidins, which are known to be regulated by SaeRS (as reviewed in (44)). While most superantigen-like proteins (SSL) were downregulated in the presence of *L. jensenii*, SSL1 was upregulated. SSL1 has been shown to have enhanced expression in nutrient-deficient media (45), which is consistent with the low glucose and nutrient competition occurring in our co-cultures. Evidence of nutrient competition is further substantiated by the upregulation of glucose and sugar phosphotransferase systems. Furthermore, nitrate metabolism was upregulated in these co-cultures, highlighted by increased expression of the oxygen-sensing system NreABC, and the nitrate reductase genes activated by this system. NreABC is typically used in anaerobic gene expression (46); considering the upregulation of nitrogen metabolism genes occurred in an aerobic setting, this would indicate that the presence *L. jensenii* alters environmental cues.

Why *L. jensenii* has apparently evolved a *tst* restrictive mechanism remains undetermined. *L. gasseri* and *L. jensenii* are found as the dominating CSTs in 6.3% and 5.3% of women respectively, while *L. crispatus* is the dominating CST in 26.2% of women (13). These frequencies indicate that *L. crispatus* may have adapted to the vaginal niche better than other species, which is supported by the presence of the glycogen-degrading pullulanase in its genome, but not in those of *L. gasseri* or *L. jensenii* (31, 32). However, given the trends on *S. aureus* at the transcriptional and translational levels between *L. crispatus* and *L. jensenii* this may suggest that the mechanism these two species use to limit the production of TSST-1 is conserved, but distinct from *L. gasseri*. A study examining the microbiota compositions of tampons from women with or without *tst*^+^ *S. aureus* did not find an association to specific CSTs, and none of the women sampled had *L. gasseri* or *L. jensenii* dominated CSTs (47). Therefore, it is unknown what the selective pressure to maintain a *tst* restrictive mechanism is for *L. jensenii*. It would be curious to examine which CST predominates following an episode of mTSS, as it would provide a strong indication for whether the TSST-1 limiting mechanisms used by the lactobacilli offer a competitive advantage in the vaginal niche.

*L. iners* has been implicated in the activation of innate immunity (48), as well as transitions to a state of BV (49), but it remains unclear whether *L. iners* is a beneficial or harmful member in the vaginal microbiota. Holm et al. (2023) proposed that different strains of *L. iners* may contribute to different phenotypes in the vaginal environment, which can either support stability or promote dysbiosis, and may be influenced by whether the *L. iners* strains were isolated from a healthy or dysbiotic environment (50). The strain used in this study, *L. iners* AB-1, is a clinical isolate from a healthy woman (51), however the origin of this strain does not appear to confer any protective benefit in the context of TSST-1 production.

*G. vaginalis* has been highly associated with dysbiosis and BV, and was found in the microbial compositions of 97.4% cases of BV, making it quite likely to be the etiological agent (52). A previous study which used *G. vaginalis* supernatant in a challenge assay for TSST-1 production found a 15% reduction in *tst* expression (53), which is consistent with the results of our luciferase assays (Fig. 1). In the current study, *G. vaginalis* enhanced production of TSST-1 from *S. aureus* at low concentrations of glucose (Fig. 3) which translated into increased T cell activation (Fig. 4) suggesting that dysbiotic BV may promote the development of mTSS. However, previous work also found increases in *tst* expression by *L. crispatus* supernatant (53), which contradicts our results (Fig. 1). The discrepancies here are likely due to the use of a non-specific rich media (BHI) in combination with high glucose VDM, which is not fully representative of the vaginal environment, and does not consider the knowledge of CcpA mediated repression that has since emerged (8).

Taken together, this work further reveals that the microbiota may play a critical role in the restraint of the superantigen TSST-1, yet this restriction is intimately linked to environmental cues. We have identified CSTs which may be more at risk than others if *tst*^+^ *S. aureus* is present in the microbiota, and have identified a potential probiotic candidate, *L. jensenii,* aimed at limiting mTSS. This study sets forward a model for examining contact-independent interactions of the vaginal microbiota and demonstrates the importance of using environment-mimicking media when studying gene regulation in dynamic environments.

## MATERIALS AND METHODS

### Ethics Statement

Healthy, adult human blood donors were recruited through the Microbiology and Immunology department at The University of Western Ontario. Donor blood was anonymized, and informed consent was obtained from donors. This protocol (110859) was approved by the London Health Sciences Centre Research Ethics Board (University of Western Ontario, London, ON, Canada).

### Bacterial growth conditions

A list and description of bacterial strains used in this study can be found in Table 1. *L. crispatus, L. gasseri,* and *L. jensenii* were routinely grown on de Man, Rogosa, and Sharpe (MRS) plates supplemented with 1.5% agar at 37°C in anaerobic conditions (BD GasPak™ EZ Anaerobe Container System) overnight, and cultured microaerophilically in MRS broth at 37°C, unless mentioned otherwise. *L. iners* and *G. vaginalis* were grown anaerobically on NYC Medium III (NYC III) plates supplemented with 1.5% agar at 37°C for two days, and cultured microaerophilically in NYC III broth at 37°C for 1–2 days, unless mentioned otherwise. *S. aureus* MN8 strains were grown on tryptic soy 1.5% agar (TSA) plates with or without 10 µg/ml chloramphenicol overnight at 37°C, and routinely cultured in TSB with or without 10 µg/ml chloramphenicol overnight at 37°C (250 rpm) unless mentioned otherwise. Vaginally Defined Media (VDM) was prepared as previously described with 0.5% proteose peptone (25, 26). Glucose concentrations were adjusted to either 60mM, 0.7mM or 0mM.

**Table 1.** Descriptions of bacterial strains used in this study.

| Strain | Description | Source |
| --- | --- | --- |
| *S. aureus* MN8 | Prototypic mTSS strain | (54) |
| *S. aureus* MN8 $\Delta ccpA$ | MN8 with a deletion of *ccpA* | (8) |
| *S. aureus* MN8 pAmilux::*P<sub>tst</sub>* | MN8 containing pAmilux::*P<sub>tst</sub>* and<br>Cm10 resistance | (5) |
| *S. aureus* MN8 $\Delta ccpA$<br>pAmilux::*P<sub>tst</sub>* | MN8 $\Delta ccpA$ containing<br>pAmilux::*P<sub>tst</sub>* and Cm10 resistance | (8) |
| *L. crispatus* ATCC 33820 | Representative strain of CST I | ATCC |
| *L. gasseri* ATCC 33323 | Representative strain of CST II | ATCC |
| *L. jensenii* ATCC 25258 | Representative strain of CST V | ATCC |
| *L. iners* AB-1 | Representative strain of CST III | (51) |
| *G. vaginalis* ATCC 14018 | Representative strain of CST IV | ATCC |

### Supernatant collection and processing

*L. crispatus*, *L. gasseri*, and *L. jensenii* were subcultured microaerophilically at 1% in VDM for overnight growth at 37°C, while *L. iners* and *G. vaginalis* were subcultured in the same conditions followed by 48 hours of growth at 37°C. This was repeated for all modified formulations of VDM. Unless otherwise mentioned, supernatants were collected by centrifuging cultures of the five representative CST strains or *S. aureus* strains at 3750 x *g* for 7 minutes (4°C). CST supernatants were filter sterilized with a 0.2 µm syringe filter (Basix) and stored at 4°C, while *S. aureus* supernatants were filter sterilized and stored at −20°C until ready for processing.

### Luciferase reporter assay

Representative CST supernatants were diluted 1/4 with their respective fresh VDM. The dilutions were used to account for changes in pH that would not sustain *S. aureus* growth, the presence of potential inhibitory molecules, as well as the depletion of nutrients from the media. Previously constructed *S. aureus* MN8 containing pAmilux::P*_tst_* and MN8 Δ*ccpA* containing pAmilux::P*_tst_* strains (Table 1) were subcultured at 1% in various formulations of VDM and each of the diluted CST supernatants, and incubated for 3 hours. The cultures were brought to an OD_600_ of 0.01 and inoculated in a 96-well plate in triplicate for 18 hours at 37°C in a Biotek Synergy H4 multimode plate reader, with continuous shaking on the medium setting. Measures of bacterial growth (OD_600_) and activity of the *tst* promoter luminescence production (RLU) were recorded once every hour. The relative expression was calculated as the area under the curve (AUC) of RLU over the AUC of OD_600_.

### Supernatant experiments

*S. aureus* MN8 and MN8 Δ*ccpA* were subcultured at 1% in various forms of VDM including the media with diluted CST supernatants. Following 3 hours, the cells were adjusted to an OD_600_ of 0.01 in the same media and grown overnight. The following day, OD_600_ readings were recorded and supernatant was precipated by a concentration of 6% trichloroacetic acid (TCA). The precipitates were washed with ice-cold acetone followed by centrifugation 15,000 x g at 4°C. The pellet was resuspended in 8M urea and stored at −20°C until usage for SDS-PAGE.

The Western blot was performed as previously described (8). Briefly, 12% acrylamide gels were run at 80-85V for 30 minutes, then 150V for 1 hour. The gel was transferred onto a polyvinylidene difluoride (PVDF) membrane and blocked overnight at 4°C with 5% skimmed milk, 10% horse serum (Gibco) and 10% fetal calf serum (Wisent) in Phosphate Buffered Saline (PBS). It was then washed and incubated for 1 hour with 1:1000 of rabbit polyclonal anti-TSST-1 antibody (55). IRDye 800-conjugated donkey anti-rabbit IgG antibody (Rockland) was added in a 1:20,000 dilution, and applied to the membrane, with incubation for 1 hour in the dark. The membrane was imaged using an Odyssey imager (LI-COR Biosciences).

### Co-cultures

To investigate the contact-independent regulation occurring between *S. aureus* and each of the protective CST species, co-cultures were performed in various formulations of VDM. *L. crispatus*, *L. gasseri*, and *L. jensenii* were grown overnight at 37°C in MRS broth. *S. aureus* MN8 and MN8 Δ*ccpA* were grown overnight at 37°C in TSB. The co-culture apparatus was prepared with a sterile 0.65 µm nylon membrane (GVS North America). Each side contained 20 mL of VDM, and a 1/100 subculture of one CST species or one of the *S. aureus* strains. The co-cultures were grown overnight at 37°C in a shaking incubator (250 rpm) to mimic the aerobic conditions of mTSS, and in a stand-still incubator to mimic low oxygen conditions otherwise seen in the vagina. In total, co-cultures were performed at 60mM, 0.7mM and 0mM glucose, as well as aerobically and microaerophilically. A diagram of the co-culture set-up can be found in Fig. S3

The following day, a TCA precipitation was performed as previously described, and untreated supernatant was collected for ELISA. TCA precipitations were stored at −20°C until use in Western blotting as previously described.

### PBMC Assay

IL-2 production from peripheral blood mononuclear cells (PBMC) was assessed by exposing the cells to *S. aureus* supernatants collected from VDM 0.7mM glucose CST supernatant experiments. The isolation of PBMCs was performed as previously described (8). PBMCs were seeded for a final concentration of 1 x 10^6^ cells/mL in a 96-well plate.

The *S. aureus* supernatants were then serially diluted and added to PBMCs in duplicate. Following an 18-hour incubation at 37°C and 5% CO_2_, the plates were centrifuged and the cell supernatants were collected for an IL-2 ELISA, using the manufacturer’s instructions (Invitrogen). The plates were read at 450nm and 570nm using a Biotek Synergy H4 multimode plate reader.

### TSST-1 ELISA

To quantify TSST-1, co-cultures of interest were reproduced three times. TSST-1 production from co-cultures of interest was determined through a sandwich ELISA using the manufacturer’s instructions (Kimberly-Clark). Briefly, supernatants from the *S. aureus* side of co-cultures were collected, filter-sterilized and stored at −20°C prior to the assay. Plates were coated with anti-TSST-1 polyclonal rabbit IgG solution, followed by addition of the samples. Detection was performed using an Anti-TSST-1 IgG Horseradish Peroxidase Conjugate. The plates were read at 450nm and 570nm using a Biotek Synergy H4 multimode plate reader.

### RNA-seq

Co-cultures of interest were incubated for 4 hours each (OD_600_ approximately 1) prior to RNA extraction. The *S. aureus* sides of the co-cultures were centrifuged and pelleted, followed by re-suspension in RNAprotect (QIAGEN), and pelleted again. Pellets were lysed using 100 µg/mL lysostaphin and RNA was extracted with the RNeasy Plus Mini Kit (QIAGEN). Extracted RNA was treated with the Turbo DNA-free Kit (Ambion) and the RNA quality was determined using an Agilent Bioanalyzer (Robarts Institute, University of Western Ontario), and only samples with RNA integrity numbers above 9 were sequenced. RNA sequencing as well as comparative analysis was performed by SeqCenter (Pittsburgh, USA).12 million paired-end Illumina sequencing was performed, followed by analysis as previously described (8). Sequences were compared against the publicly available MN8 genome (GenBank assembly accession no. GCA_024296845.1).

### Statistical Analysis

All statistical analysis was performed using GraphPad Prism 9. Ordinary one-way ANOVA was used without correction for multiple comparisons for luciferase assays and IL-2 ELISAs. Unpaired t-tests were used for TSST-1 ELISAs.

### Data Availability

Raw RNA read data were deposited at NCBI under BioProject accession no. PRJNA991630.

## REFERENCES

1. Tuffs SW, Haeryfar SMM, McCormick JK. 2018. Manipulation of Innate and Adaptive Immunity by Staphylococcal Superantigens. Pathogens 7:E53.

2. Sundberg EJ, Deng L, Mariuzza RA. 2007. TCR recognition of peptide/MHC class II complexes and superantigens. Semin Immunol 19:262–271.

3. CDC. 2021. Toxic Shock Syndrome (Other Than Streptococcal) (TSS) 2011 Case Definition | CDC. https://ndc.services.cdc.gov/case-definitions/toxic-shock-syndrome-2011/. Retrieved 30 May 2022.

4. Schlievert PM, Davis CC. 2020. Device-Associated Menstrual Toxic Shock Syndrome. Clinical Microbiology Reviews 33:e00032–19.

5. Baroja ML, Herfst CA, Kasper KJ, Xu SX, Gillett DA, Li J, Reid G, McCormick JK. 2016. The SaeRS Two-Component System Is a Direct and Dominant Transcriptional Activator of Toxic Shock Syndrome Toxin 1 in *Staphylococcus aureus*. J Bacteriol 198:2732–2742.

6. Tuffs SW, Herfst CA, Baroja ML, Podskalniy VA, DeJong EN, Coleman CEM, McCormick JK. 2019. Regulation of toxic shock syndrome toxin-1 by the accessory gene regulator in *Staphylococcus aureus* is mediated by the repressor of toxins. Mol Microbiol 112:1163– 1177.

7. Seidl K, Bischoff M, Berger-Bächi B. 2008. CcpA mediates the catabolite repression of *tst* in *Staphylococcus aureus*. Infect Immun 76:5093–5099.

8. Dufresne K, Podskalniy VA, Herfst CA, Lovell GFM, Lee IS, DeJong EN, McCormick JK, Tuffs SW. 2022. Glucose Mediates Niche-Specific Repression of *Staphylococcus aureus* Toxic Shock Syndrome Toxin-1 through the Activity of CcpA in the Vaginal Environment. Journal of Bacteriology 0:e00269–22.

9. Yarwood JM, McCormick JK, Schlievert PM. 2001. Identification of a Novel Two-Component Regulatory System That Acts in Global Regulation of Virulence Factors of *Staphylococcus aureus*. Journal of Bacteriology 183:1113–1123.

10. Pragman AA, Ji Y, Schlievert PM. 2007. Repression of *Staphylococcus aureus* SrrAB using inducible antisense *srrA* alters growth and virulence factor transcript levels. Biochemistry 46:314–321.

11. McCormick JK, Yarwood JM, Schlievert PM. 2001. Toxic shock syndrome and bacterial superantigens: an update. Annu Rev Microbiol 55:77–104.

12. Kehrberg MW, Latham RH, Haslam BT, Hightower A, Tanner M, Jacobson JA, Barbour AG, Noble V, Smith CB. 1981. Risk factors for staphylococcal toxic-shock syndrome. Am J Epidemiol 114:873–879.

13. Ravel J, Gajer P, Abdo Z, Schneider GM, Koenig SSK, McCulle SL, Karlebach S, Gorle R, Russell J, Tacket CO, Brotman RM, Davis CC, Ault K, Peralta L, Forney LJ. 2011. Vaginal microbiome of reproductive-age women. Proc Natl Acad Sci U S A 108 Suppl 1:4680–4687.

14. Chee WJY, Chew SY, Than LTL. 2020. Vaginal microbiota and the potential of *Lactobacillus* derivatives in maintaining vaginal health. Microb Cell Fact 19:203.

15. Gajer P, Brotman RM, Bai G, Sakamoto J, Schütte UME, Zhong X, Koenig SSK, Fu L, Ma ZS, Zhou X, Abdo Z, Forney LJ, Ravel J. 2012. Temporal dynamics of the human vaginal microbiota. Sci Transl Med 4:132ra52.

16. Onderdonk AB, Delaney ML, Fichorova RN. 2016. The Human Microbiome during Bacterial Vaginosis. Clin Microbiol Rev 29:223–238.

17. Hillier SL, Lau RJ. 1997. Vaginal microflora in postmenopausal women who have not received estrogen replacement therapy. Clin Infect Dis 25 Suppl 2:S123–126.

18. Amabebe E, Anumba DOC. 2018. The Vaginal Microenvironment: The Physiologic Role of Lactobacilli. Front Med (Lausanne) 5:181.

19. Krog MC, Hugerth LW, Fransson E, Bashir Z, Nyboe Andersen A, Edfeldt G, Engstrand L, Schuppe-Koistinen I, Nielsen HS. 2022. The healthy female microbiome across body sites: effect of hormonal contraceptives and the menstrual cycle. Hum Reprod 37:1525–1543.

20. Gliniewicz K, Schneider GM, Ridenhour BJ, Williams CJ, Song Y, Farage MA, Miller K, Forney LJ. 2019. Comparison of the Vaginal Microbiomes of Premenopausal and Postmenopausal Women. Frontiers in Microbiology 10.

21. Winston McPherson G, Long T, Salipante SJ, Rongitsch JA, Hoffman NG, Stephens K, Penewit K, Greene DN. 2019. The Vaginal Microbiome of Transgender Men. Clin Chem 65:199–207.

22. Mirmonsef P, Hotton AL, Gilbert D, Gioia CJ, Maric D, Hope TJ, Landay AL, Spear GT. 2016. Glycogen Levels in Undiluted Genital Fluid and Their Relationship to Vaginal pH, Estrogen, and Progesterone. PLoS One 11:e0153553.

23. Mirmonsef P, Hotton AL, Gilbert D, Burgad D, Landay A, Weber KM, Cohen M, Ravel J, Spear GT. 2014. Free Glycogen in Vaginal Fluids Is Associated with *Lactobacillus* Colonization and Low Vaginal pH. PLoS One 9:e102467.

24. Mirmonsef P, Modur S, Burgad D, Gilbert D, Golub ET, French AL, McCotter K, Landay AL, Spear GT. 2015. An exploratory comparison of vaginal glycogen and *Lactobacillus* levels in pre- and post-menopausal women. Menopause 22:702–709.

25. Geshnizgani AM, Onderdonk AB. 1992. Defined medium simulating genital tract secretions for growth of vaginal microflora. J Clin Microbiol 30:1323–1326.

26. Anukam KC, Reid G. 2008. Effects of metronidazole on growth of *Gardnerella vaginalis* ATCC 14018, probiotic *Lactobacillus rhamnosus* GR-1 and vaginal isolate *Lactobacillus plantarum* KCA. Microbial Ecology in Health and Disease 20:48–52.

27. Regassa LB, Novick RP, Betley MJ. 1992. Glucose and nonmaintained pH decrease expression of the accessory gene regulator (*agr*) in *Staphylococcus aureus*. Infect Immun 60:3381–3388.

28. Regassa LB, Betley MJ. 1992. Alkaline pH decreases expression of the accessory gene regulator (*agr*) in *Staphylococcus aureus*. J Bacteriol 174:5095–5100.

29. Nasioudis D, Beghini J, Bongiovanni AM, Giraldo PC, Linhares IM, Witkin SS. 2015. α-Amylase in Vaginal Fluid: Association With Conditions Favorable to Dominance of *Lactobacillus*. Reprod Sci 22:1393–1398.

30. Nunn KL, Clair GC, Adkins JN, Engbrecht K, Fillmore T, Forney LJ. 2020. Amylases in the Human Vagina. mSphere 5:e00943–20.

31. Zhang J, Li L, Zhang T, Zhong J. 2022. Characterization of a novel type of glycogen-degrading amylopullulanase from *Lactobacillus crispatus*. Appl Microbiol Biotechnol https://doi.org/10.1007/s00253-022-11975-2.

32. Woolston BM, Jenkins DJ, Hood-Pishchany MI, Nahoum SR, Balskus EP. 2021. Characterization of vaginal microbial enzymes identifies amylopullulanases that support growth of *Lactobacillus crispatus* on glycogen. bioRxiv https://doi.org/10.1101/2021.07.19.452977.

33. Spear GT, French AL, Gilbert D, Zariffard MR, Mirmonsef P, Sullivan TH, Spear WW, Landay A, Micci S, Lee B-H, Hamaker BR. 2014. Human α-amylase present in lower-genital-tract mucosal fluid processes glycogen to support vaginal colonization by *Lactobacillus*. J Infect Dis 210:1019–1028.

34. Spear GT, McKenna M, Landay AL, Makinde H, Hamaker B, French AL, Lee B-H. 2015. Effect of pH on Cleavage of Glycogen by Vaginal Enzymes. PLoS One 10:e0132646.

35. Lakshmi HP, Prasad UV, Yeswanth S, Swarupa V, Prasad OH, Narasu ML, Sarma PVGK. 2013. Molecular characterization of α-amylase from *Staphylococcus aureus*. Bioinformation 9:281–285.

36. O’Hanlon DE, Moench TR, Cone RA. 2011. In vaginal fluid, bacteria associated with bacterial vaginosis can be suppressed with lactic acid but not hydrogen peroxide. BMC Infect Dis 11:200.

37. Gong Z, Luna Y, Yu P, Fan H. 2014. Lactobacilli Inactivate *Chlamydia trachomatis* through Lactic Acid but Not H2O2. PLOS ONE 9:e107758.

38. Delgado-Diaz DJ, Jesaveluk B, Hayward JA, Tyssen D, Alisoltani A, Potgieter M, Bell L, Ross E, Iranzadeh A, Allali I, Dabee S, Barnabas S, Gamieldien H, Blackburn JM, Mulder N, Smith SB, Edwards VL, Burgener AD, Bekker L-G, Ravel J, Passmore J-AS, Masson L, Hearps AC, Tachedjian G. 2022. Lactic acid from vaginal microbiota enhances cervicovaginal epithelial barrier integrity by promoting tight junction protein expression. Microbiome 10:141.

39. Schlievert PM, Gaitán AV, Kilgore SH, Roe AL, Maukonen J, Lehtoranta L, Leung DYM, Marsman DS. 2023. Inhibition of Toxic Shock Syndrome-Associated *Staphylococcus aureus* by Probiotic Lactobacilli. Microbiol Spectr e0173523.

40. Witkin SS, H M-S, Im L, A J, Wj L, Lj F. 2013. Influence of vaginal bacteria and D- and L-lactic acid isomers on vaginal extracellular matrix metalloproteinase inducer: implications for protection against upper genital tract infections. mBio 4.

41. Beghini J, Linhares IM, Giraldo PC, Ledger WJ, Witkin SS. 2015. Differential expression of lactic acid isomers, extracellular matrix metalloproteinase inducer, and matrix metalloproteinase-8 in vaginal fluid from women with vaginal disorders. BJOG 122:1580– 1585.

42. Li J, Wang W, Xu SX, Magarvey NA, McCormick JK. 2011. *Lactobacillus reuteri*-produced cyclic dipeptides quench *agr*-mediated expression of toxic shock syndrome toxin-1 in staphylococci. Proc Natl Acad Sci U S A 108:3360–3365.

43. Younes JA, Reid G, van der Mei HC, Busscher HJ. 2016. Lactobacilli require physical contact to reduce staphylococcal TSST-1 secretion and vaginal epithelial inflammatory response. Pathog Dis 74:ftw029.

44. Liu Q, Yeo W-S, Bae T. 2016. The SaeRS Two-Component System of *Staphylococcus aureus*. Genes (Basel) 7:E81.

45. Bretl DJ, Elfessi A, Watkins H, Schwan WR. 2019. Regulation of the Staphylococcal Superantigen-Like Protein 1 Gene of Community-Associated Methicillin-Resistant *Staphylococcus aureus* in Murine Abscesses. Toxins (Basel) 11:391.

46. Schlag S, Fuchs S, Nerz C, Gaupp R, Engelmann S, Liebeke M, Lalk M, Hecker M, Götz F. 2008. Characterization of the oxygen-responsive NreABC regulon of *Staphylococcus aureus*. J Bacteriol 190:7847–7858.

47. Jacquemond I, Muggeo A, Lamblin G, Tristan A, Gillet Y, Bolze PA, Bes M, Gustave CA, Rasigade J-P, Golfier F, Ferry T, Dubost A, Abrouk D, Barreto S, Prigent-Combaret C, Thioulouse J, Lina G, Muller D. 2018. Complex ecological interactions of *Staphylococcus aureus* in tampons during menstruation. Sci Rep 8:9942.

48. Doerflinger SY, Throop AL, Herbst-Kralovetz MM. 2014. Bacteria in the vaginal microbiome alter the innate immune response and barrier properties of the human vaginal epithelia in a species-specific manner. J Infect Dis 209:1989–1999.

49. Tamarelle J, Shardell MD, Ravel J, Brotman RM. 2022. Factors Associated With Incidence and Spontaneous Clearance of Molecular-Bacterial Vaginosis: Results From a Longitudinal Frequent-Sampling Observational Study. Sex Transm Dis 49:649–656.

50. Holm JB, Carter KA, Ravel J, Brotman RM. 2023. *Lactobacillus iners* and genital health: molecular clues to an enigmatic vaginal species. Curr Infect Dis Rep 25:67–75.

51. Macklaim JM, Gloor GB, Anukam KC, Cribby S, Reid G. 2011. At the crossroads of vaginal health and disease, the genome sequence of *Lactobacillus iners* AB-1. Proc Natl Acad Sci U S A 108:4688–4695.

52. Srinivasan S, Hoffman NG, Morgan MT, Matsen FA, Fiedler TL, Hall RW, Ross FJ, McCoy CO, Bumgarner R, Marrazzo JM, Fredricks DN. 2012. Bacterial communities in women with bacterial vaginosis: high resolution phylogenetic analyses reveal relationships of microbiota to clinical criteria. PLoS One 7:e37818.

53. MacPhee RA, Miller WL, Gloor GB, McCormick JK, Hammond J-A, Burton JP, Reid G. 2013. Influence of the vaginal microbiota on toxic shock syndrome toxin 1 production by *Staphylococcus aureus*. Appl Environ Microbiol 79:1835–1842.

54. Blomster-Hautamaa DA, Kreiswirth BN, Kornblum JS, Novick RP, Schlievert PM. 1986. The nucleotide and partial amino acid sequence of toxic shock syndrome toxin-1. J Biol Chem 261:15783–15786.

55. Schlievert PM, Blomster DA. 1983. Production of staphylococcal pyrogenic exotoxin type C: influence of physical and chemical factors. J Infect Dis 147:236–242.

