## Supplemental figures for "Vaginal Community State Types (CSTs) Alter Environmental Cues and Production of the *Staphylococcus aureus* Toxic Shock Syndrome Toxin-1 (TSST-1)"

Running Title: Vaginal CSTs alter production of TSST-1

Keywords: *Staphylococcus aureus*, TSST-1, glucose, toxic shock syndrome, mTSS, vaginal microbiota, lactobacilli, *Gardnerella vaginalis*

**SUPPLEMENTAL FIGURES**

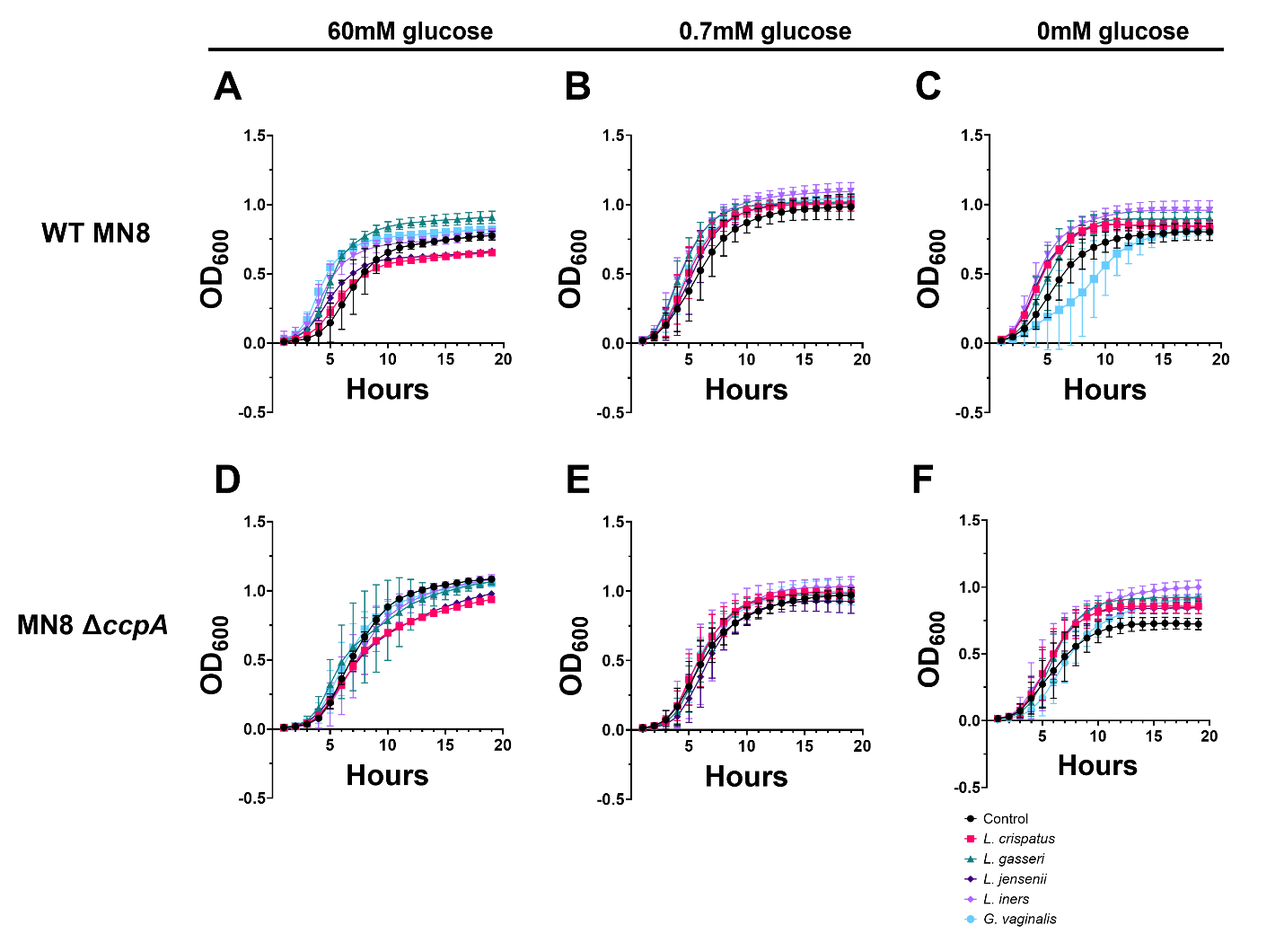
**Fig. S1 Growth curves for *S. aureus* grown in CST supernatants.** CSTs I-V were grown in standard VDM (**A, D**), or modified VDM: 0.7mM glucose (**B, E**), 0mM glucose (**C, F**). The supernatants were sterilized and diluted with the respective fresh media at a 1/4 dilution. Wild-type MN8 and MN8 ∆*ccpA* containing the luminescence reporter plasmid pAmilux::P*_tst_* were then grown in the diluted supernatants, and curves were adjusted to represent start of growth relative to fresh media. The assay was performed in the Synergy H4 reader at 37°C and continuous shaking on the medium setting.

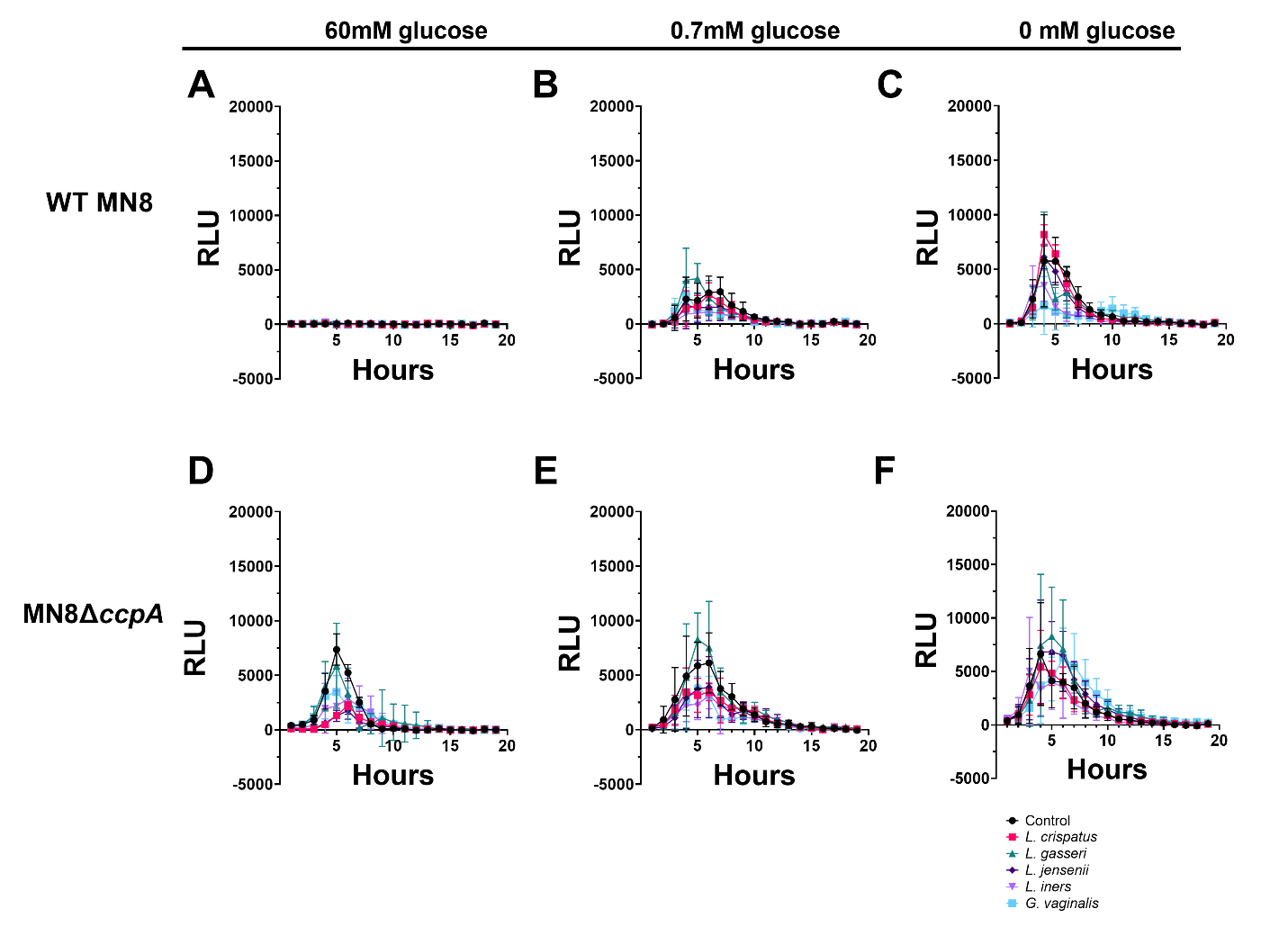
**Fig. S2 Luminescence curves for *S. aureus* grown in CST supernatants.** CSTs I-V were grown in standard VDM (**A, D**), or modified VDM: 0.7mM glucose (**B, E**), 0mM glucose (**C, F**). The supernatants were sterilized and diluted with the respective fresh media at a 1/4 dilution. Wild-type MN8 and MN8 ∆*ccpA* containing the luminescence reporter plasmid pAmilux::P*_tst_* were then grown in the diluted supernatants, and luminescence curves were adjusted to represent start of growth relative to fresh media. The assay was performed in the Synergy H4 reader at 37°C and continuous shaking on the medium setting.

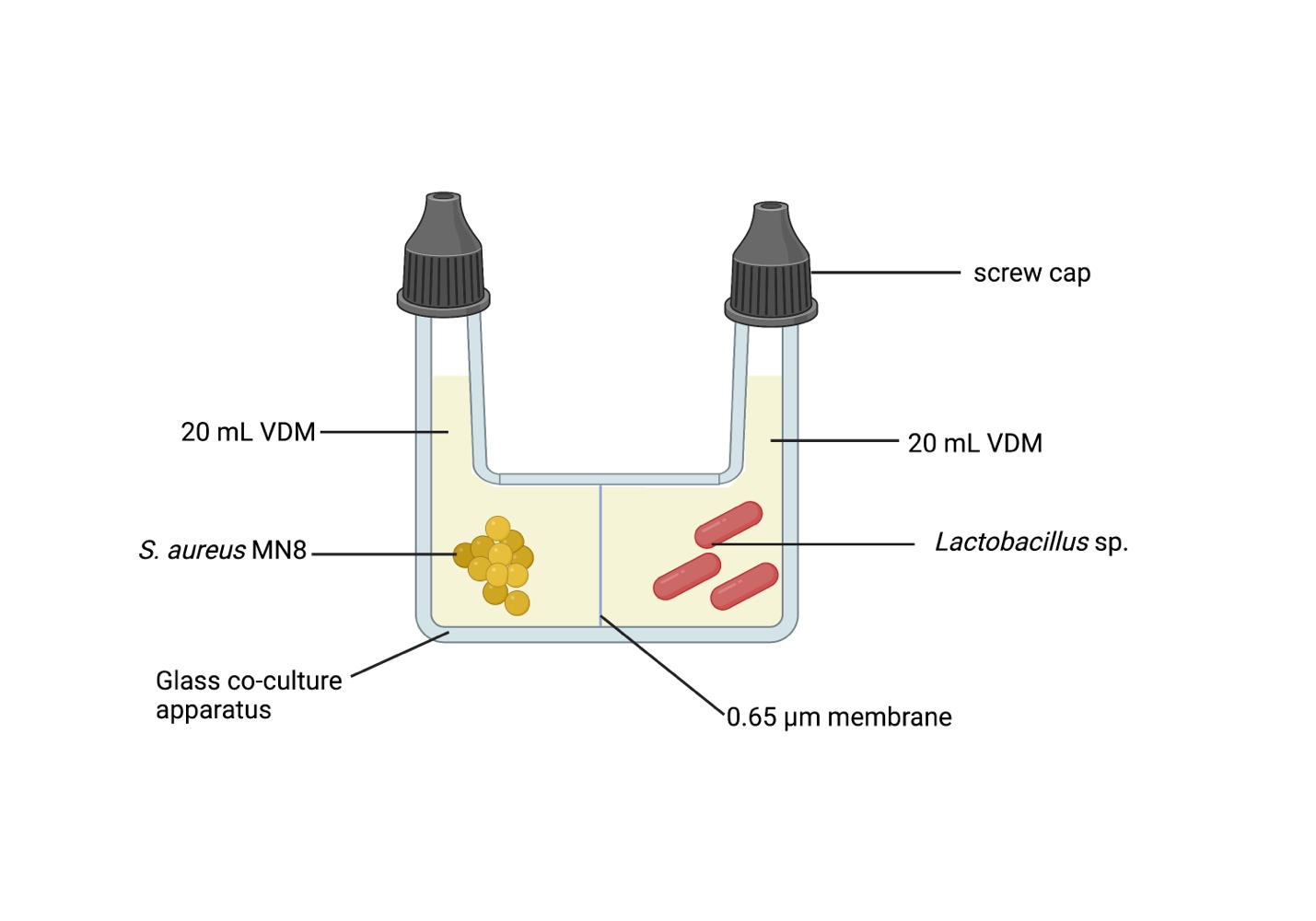

**Fig. S3 Co-cultures apparatus setup.** A glass co-culture apparatus was washed and autoclaved prior to set up. Two sterile O-rings were placed on either side of a 0.65 µm membrane, and the whole apparatus was re-enforced with a sterile clamp. To either side of the membrane, 20 mL of VDM was added, followed by 200 µl of overnight bacterial cultures. Control co-cultures were performed with the same *S. aureus* strain on either side of the membrane. Experimental co-cultures were performed with one *S. aureus* strain on the left, and one of three *Lactobacillus* spp. on the right. Co-cultures were incubated at 37°C either with 250 rpm to mimic the aeration present in mTSS, or with no rotations to mimic the microaerophilic/anaerobic conditions of healthy menstruation. Samples were collected exclusively from the *S. aureus* side for further experimentation (TCA precipitations, TSST-1 ELISAs). Image created with biorender.com.

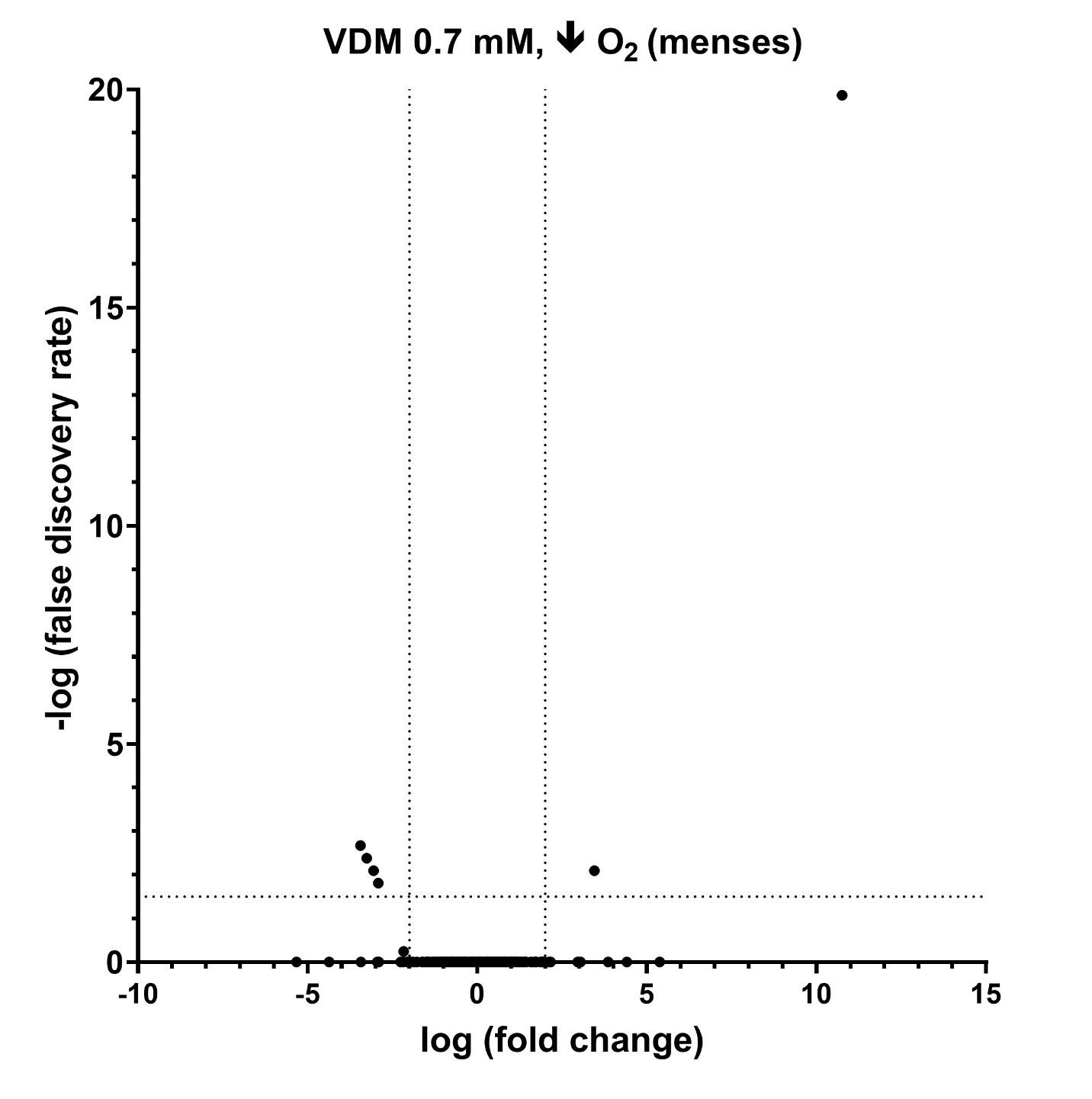

**Fig. S4 *L. jensenii* in co-culture with *S. aureus* MN8 ∆*ccpA* does not alter gene expression in conditions mimicking menses.** A co-culture apparatus separated MN8 ∆*ccpA* from *L. jensenii* with a 0.65 µm filter, with each side containing 20 mL of VDM 0.7mM glucose. Control co-cultures were also performed with MN8 ∆*ccpA* on either side of the membrane. Co-cultures were performed microaerophilically without shaking at 37°C to mimic healthy menstruation. Each dot in the volcano plot represents a single transcript, and those above the horizontal threshold indicate P < 0.001, while vertical thresholds indicate log fold changes greater than 2 in the presence of *L. jensenii*.

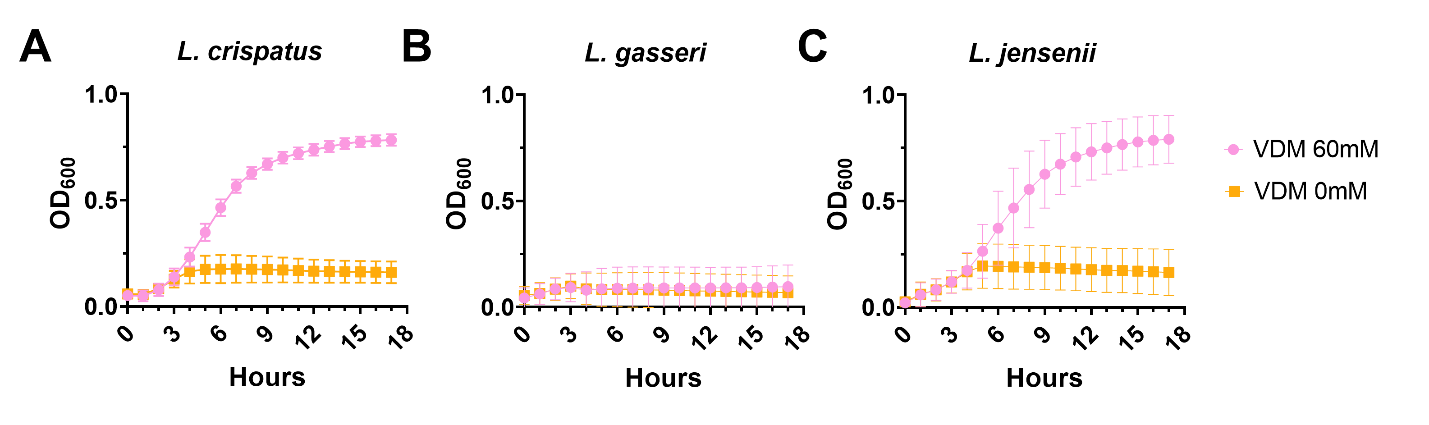
**Fig. S5 Lactobacilli growth curves in VDM 60mM and 0mM glucose.** Strains were grown routinely as described in the methods. The cultures were diluted to an OD_600_ of 0.05, and grown overnight at 37°C in the Biotek Synergy H4 multimode plate reader in VDM 60mM or VDM 0mM. Readings were taken hourly with shaking for 5 seconds on the medium setting prior to reads. Data points represent the means of the replications, and error bars represent SD. (**A**) *L. crispatus* (**B**) *L. gasseri* (**C**) *L. jensenii.*
